# ATM modulates nuclear mechanics by regulating lamin A levels

**DOI:** 10.1101/2022.02.28.482418

**Authors:** Pragya Shah, Connor W. McGuigan, Svea Cheng, Claire Vanpouille-Box, Sandra Demaria, Robert S. Weiss, Jan Lammerding

## Abstract

Ataxia-telangiectasia mutated (ATM) is one of the three main apical kinases at the crux of DNA damage response and repair in mammalian cells. ATM activates a cascade of downstream effector proteins to regulate DNA repair and cell cycle checkpoints in response to DNA double-strand breaks. While ATM is predominantly known for its role in DNA damage response and repair, new roles of ATM have recently begun to emerge, such as in regulating oxidative stress or metabolic pathways. Here, we report the surprising discovery that ATM inhibition and deletion lead to reduced expression of the nuclear envelope protein lamin A. Lamins are nuclear intermediate filaments that modulate nuclear shape, structure, and stiffness. Accordingly, inhibition or deletion of ATM resulted in increased nuclear deformability and enhanced cell migration through confined spaces, which requires substantial nuclear deformation. These findings point to a novel connection between ATM and lamin A and may have broad implications for cells with ATM mutations—as found in patients suffering from Ataxia Telangiectasia and many human cancers—which could lead to enhanced cell migration and increased metastatic potential.

## Introduction

ATM was first discovered in patients suffering from ataxia-telangiectasia (A-T), an autosomal-recessive genetic disease characterized by dilated blood vessels, neurodegeneration, immunodeficiency and predisposition to certain cancers (Boder and Sedgwick, 1958; Chun and Gatti, 2004; Chiam et al., 2011; Reiman et al., 2011; Taylor et al., 2015). ATM, along with ATM and Rad3 related (ATR) and DNA-dependent protein kinase (DNA-PK), was later identified as one of the three main apical kinases involved in the DNA damage response (DDR) pathway that are important to repair DNA double-strand breaks (Bensimon et al., 2011; Shiloh and Ziv, 2013; Awasthi et al., 2015; Blackford and Jackson, 2017). ATM, ATR, and DNA-PK belong to the PI3K-related kinase (PIKK) family of proteins, sharing structural similarities in their kinase domain as well as substrate specificity for phosphorylation (Perry and Kleckner, 2003; Lempiainen and Halazonetis, 2009; Lovejoy and Cortez, 2009; Awasthi et al., 2015; Blackford and Jackson, 2017). Upon DNA double-strand break induction, these kinases phosphorylate and regulate a cascade of downstream effector proteins to mediate DNA double-strand break repair, cell-cycle regulation, and apoptosis to minimize the risk of propagating genomically unstable cells (Shiloh, 2006; Matsuoka et al., 2007; Cheng and Chen, 2010; Smith et al., 2010; Bensimon et al., 2011; Awasthi et al., 2015; Blackford and Jackson, 2017).

While ATM is predominantly known for its contribution to DNA double-strand break repair, recent studies have implicated ATM in oxidative stress responses (Guo et al., 2010a; Guo et al., 2010b; Cosentino et al., 2011; Ditch and Paull, 2012) and various metabolic processes (Schneider et al., 2006; Li et al., 2009; Alexander and Walker, 2010; Yang et al., 2011; Daugherity et al., 2012; Ditch and Paull, 2012), including insulin signalling (Viniegra et al., 2005; Espach et al., 2015), Akt activation (Halaby et al., 2008; Li and Yang, 2010), activation of the AMPK pathway (Suzuki et al., 2004; Sun et al., 2007; Fu et al., 2008; Alexander et al., 2010), and NF-ĸB pathways (Hinz et al., 2010; Miyamoto, 2011), suggesting that ATM has various functions outside of DDR pathways.

In this study, we investigated the crosstalk between ATM and lamins. Lamins are intermediate filaments underlying the inner nuclear membrane (Gerace et al., 1978; Gerace and Huber, 2012). Lamins can be grouped into two types: A-type lamins (lamins A and C, produced by alternative splicing of the *LMNA* gene) and B-type lamins (lamins B1 and B2, encoded by the *LMNB1* and *LMNB2* genes, respectively) (Gerace et al., 1978), which play a wide variety of roles including gene regulation, cytoskeletal organization, nuclear stiffness modulation, differentiation and cell metabolism (Ho and Lammerding, 2012; Schreiber and Kennedy, 2013; Davidson et al., 2014; Gruenbaum and Foisner, 2015; Bell and Lammerding, 2016; Xie and Burke, 2016; Charar and Gruenbaum, 2017). Mutations in lamins, particularly in the *LMNA* gene, can cause a broad range of diseases such as Hutchinson-Gilford Progeria Syndrome (HGPS), Emery-Dreifuss Muscular Dystrophy, and autosomal dominant leukodystrophy, collectively referred to as laminopathies (Charar and Gruenbaum, 2017; Wong and Stewart, 2020). Previous studies have shown that ATM is mislocalized in patients suffering from HGPS or restricted dermopathy, a disease caused by impaired lamin A processing (Liu et al., 2006; Dechat et al., 2008). ATR has also been shown to mislocalize in HeLa cells expressing *LMNA* mutations (Manju et al., 2006), suggesting a crosstalk between DDR kinases and A-type lamins. Although these previous studies have established a role of A-type lamins in modulating ATM, ATR, and DDR pathways (Gonzalo, 2014), it remains unclear whether DDR kinases, and ATM in particular, can conversely affect lamin A/C levels or function.

In this work, we used chemical inhibitors, along with either stable deletion or inducible depletion of ATM to study the effect of ATM on lamins. We found that functional loss of ATM reduced lamin A protein levels across multiple cell lines. This effect was specific to ATM, as inhibition of ATR or DNA-PK did not cause significant alterations to lamin A protein levels. Moreover, inhibition or depletion of ATM resulted in increased nuclear deformability and enhanced cell migration speed through confined spaces, consistent with reduced lamin A expression. Although the exact mechanism underlying ATM’s regulation of lamin A remains to be elucidated, we identified that ATM inhibition or deletion led to reduced *LMNA* mRNA levels, suggesting altered transcriptional regulation as a potential candidate. Collectively, we have uncovered a novel role for ATM in modulating nuclear stiffness and confined migration by regulating lamin A levels.

## Results

### Inhibition of ATM but not DNA-PK or ATR reduces lamin A protein levels

To study the impact of specific DDR kinase activity on lamin levels, we selectively inhibited ATM, ATR and DNA-PK using small-molecule chemical inhibitors in two independent human cell lines— a breast cancer cell line (MDA-MB-231) and a fibrosarcoma cell line (HT1080)— and assessed lamin A/C protein levels after 48 hours of treatment by Western analysis. ATM inhibition using KU-55933 led to significant reduction in lamin A protein levels compared to vehicle-treated controls cells in both MDA-MB-231 and HT1080 cells (Fig. 1A-C). Lamin C, which is encoded by the same *LMNA* gene, showed a similar trend, but the reduction was not statistically significant (Fig. 1A-C). Interestingly, lamin B1 levels were not reduced following ATM inhibition (Suppl. Fig. 1A-B). In contrast to ATM inhibition, pharmacological inhibition of ATR (Fig. 1D-F) or DNA-PK (Fig. 1G-I) using VE-821 and NU7441, respectively, did not significantly alter lamin A levels, consistent with a previous study that found that although depletion of ATR affected nuclear shape and stiffness, it did not alter lamin A/C levels (Kidiyoor et al., 2020). Taken together, these findings suggest that ATM, but not other DDR kinases, regulates lamin A levels.

**Figure 1:**
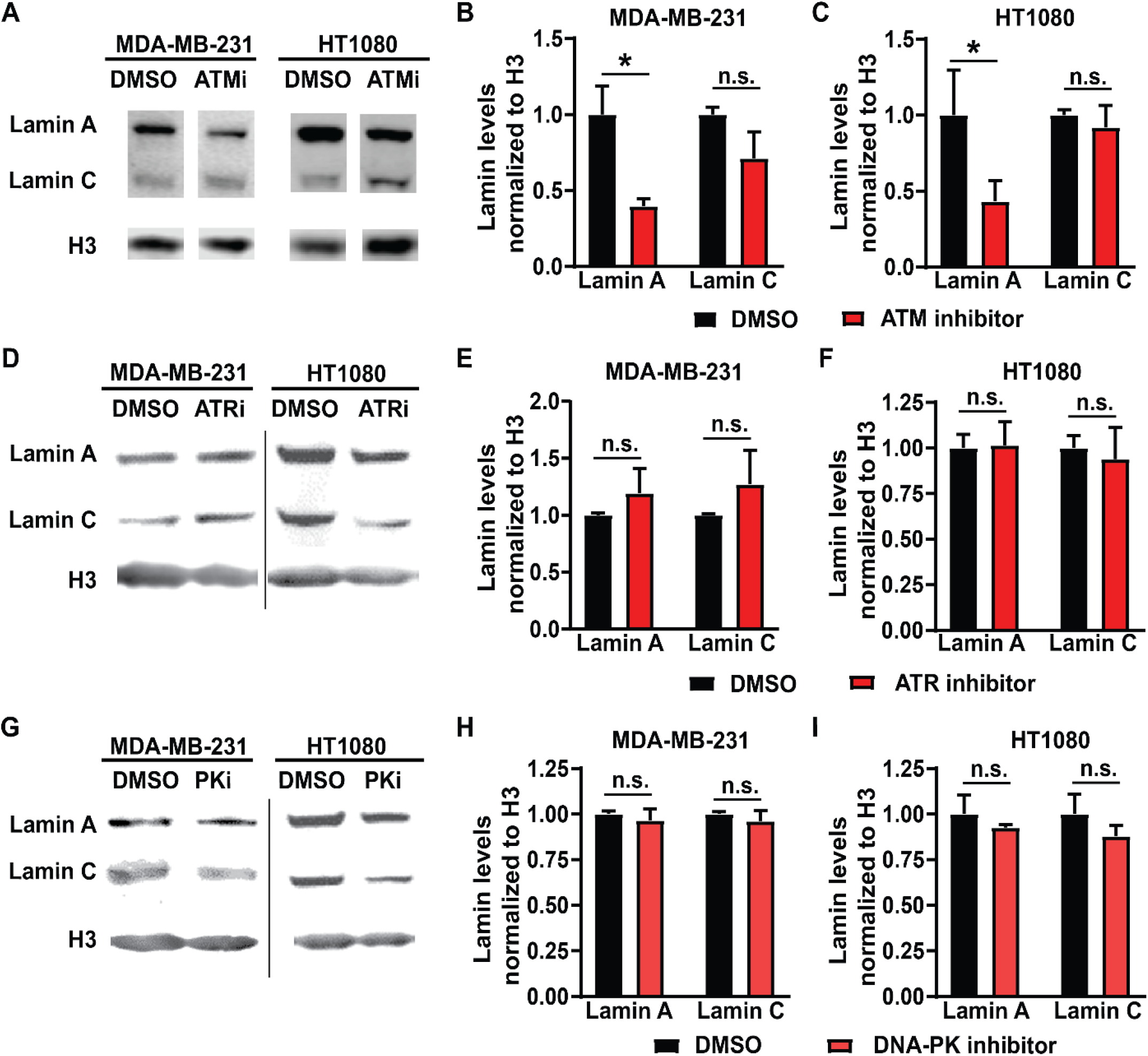
Inhibition of ATM but not ATR or DNA-PK reduces lamin A levels. **(A)** Representative immunoblot for lamin A and lamin C of MDA-MB-231 and HT1080 cells treated with 10 µM of ATM inhibitor KU-55933 or 0.1% DMSO for 48 hours. Histone H3 was used as loading control. **(B)** Quantification of lamin A and C protein levels from immunoblots of MDA-MB-231 cells, based on three independent experiments. Lamin levels were normalized to DMSO treated cells. *, *p* < 0.05 based on Two-way ANOVA with Tukey’s multiple comparison test. *N* = 3. **(C)** Quantification of lamin A and C protein levels from immunoblots of HT1080 cells, based on three independent experiments. Lamin levels were normalized to DMSO treated cells. *, *p* < 0.05 based on Two-way ANOVA with Tukey’s multiple comparison test. *N* = 3 **(D)** Representative immunoblot for lamin A and lamin C of MDA-MB-231 and HT1080 cells treated with 10 µM of ATR inhibitor VE-821 or 0.1% DMSO for 48 hours. Histone H3 was used as loading control. **(E)** Quantification of lamin A and C protein levels from immunoblots for MDA-MB-231 cells, based on three independent experiments. Lamin levels were normalized to DMSO treated cells. Differences were not statistically significant (n.s.) based on two-way ANOVA. *N* = 3. **(F)** Quantification of lamin A and C protein levels from immunoblots for HT1080 cells, based on three independent experiments. Lamin levels were normalized to DMSO treated cells. Differences were not statistically significant (n.s.) based on two-way ANOVA. *N* = 3 **(G)** Representative immunoblot for lamin A and lamin C of MDA-MB-231 and HT1080 cells treated with 5 µM of DNA-PK inhibitor NU7441 or 0.1% DMSO for 48 hours. Histone H3 was used as loading control. **(H)** Quantification of lamin A and C protein levels from immunoblots for MDA-MB-231 cells, based on three independent experiments. Lamin levels were normalized to DMSO treated cells. Differences were not statistically significant (n.s.) based on two-way ANOVA. *N* = 3. **(I)** Quantification of lamin A and C protein levels from immunoblots for HT1080 cells, based on three independent experiments. Lamin levels were normalized to DMSO treated cells. Differences were not statistically significant (n.s.) based on two-way ANOVA. *N* = 3. Error bars in this figure represent mean ± s.e.m.

### Stable or inducible knockdown of ATM also leads to low lamin A levels

Chemical inhibitors can have off-target effects or impact multiple proteins and pathways (Wynn et al., 2011). To confirm that reduction of lamin A levels was specifically mediated through ATM, we assessed lamin levels after depletion of ATM using two independent genetic approaches. Mouse embryonic fibroblasts (MEFs) in which the gene encoding ATM was deleted (‘*Atm*-Null’) (Balmus et al., 2012), along with corresponding wild-type controls, were used to assess the effect of stable deletion of *Atm* on lamin levels. To avoid potential selection bias due to long-term deletion of *Atm*, we generated 4T1 mouse mammary carcinoma cells with doxycycline inducible expression of shRNA targeting either *Atm*, leading to ATM depletion within 4 days as confirmed by immunoblot analysis, or a non-target control (Suppl. Fig. 2). Since high doses of doxycycline can induce DNA damage and DDR pathways (Song et al., 2014; Lamb et al., 2015) and to avoid potentially confounding non-specific effects of doxycycline treatment, we limited our analysis in the 4T1 cells to comparing the groups that had received identical low doses (4 μg/ml) of doxycycline. Both 4T1 cells depleted for ATM (Fig. 2A-B) and *Atm*-Null MEFs (Fig. 2C-D) had significantly reduced lamin A protein levels compared to wild-type or non-target control cells, respectively, but no impact on lamin B levels (Suppl. Fig. 1C-D) when assessed by Western analysis, similar to the effect of pharmacological ATM inhibition on lamin levels (Fig. 1A-C, Suppl. Fig. 1A-B). Deletion or depletion of *Atm* also led to a significant reduction in lamin C levels in both MEFs and 4T1 cells. To corroborate these results further, we quantified lamin A/C levels in *Atm*-Null and matching wild-type control MEFs by immunofluorescence labeling, which revealed reduced lamin A/C expression in *Atm*-Null MEFs compared to wild-type MEFs (Fig. 2E-F). Surprisingly, Atm-Null MEFs also had smaller nuclear cross-sectional areas (Fig. 2G). Collectively, these findings suggest that inhibition or deletion of ATM reduces lamin A protein levels in various human and murine cell lines.

**Figure 2:**
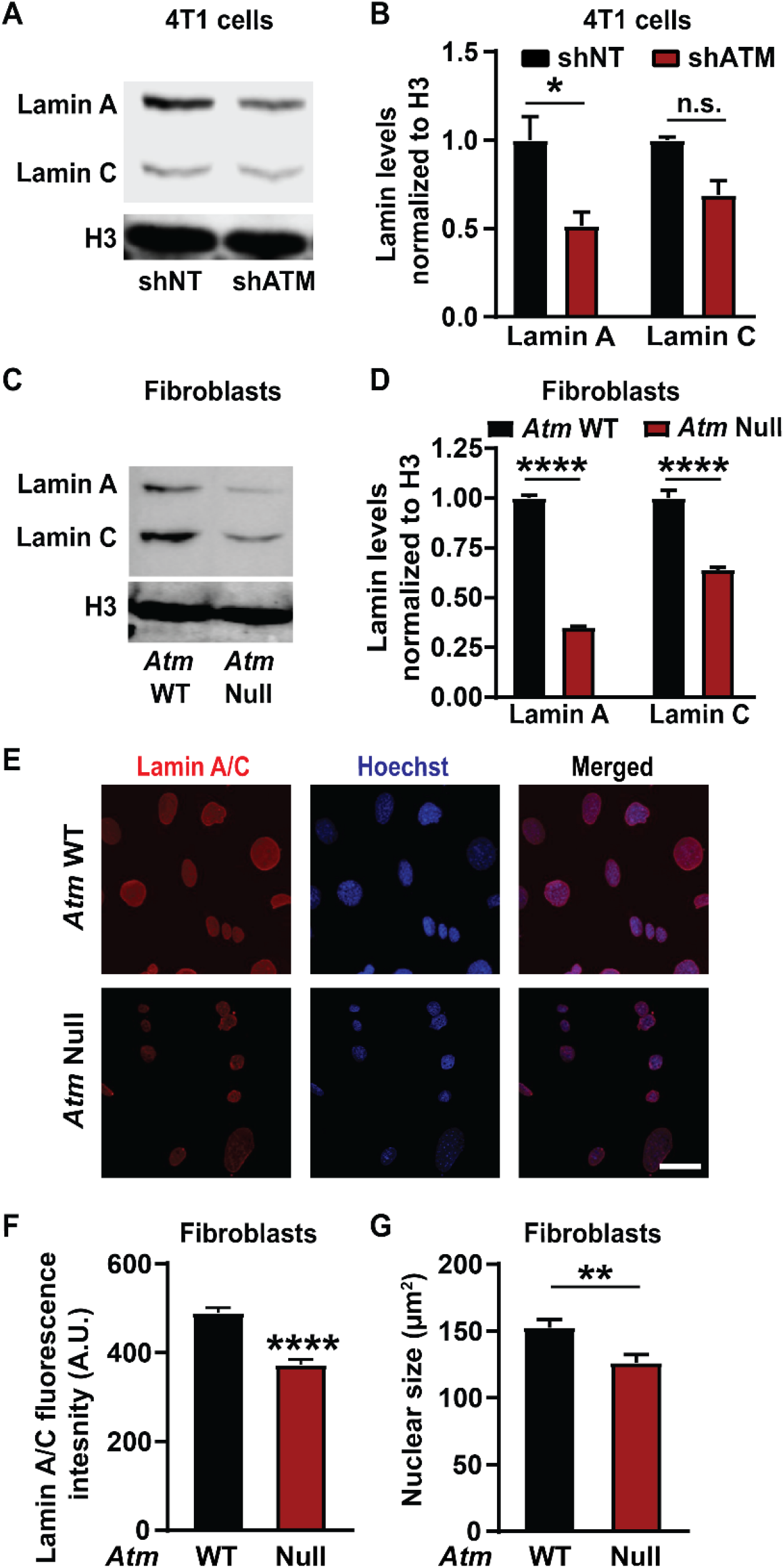
ATM deletion and depletion reduce lamin A levels. **(A)** Representative immunoblot for lamin A and lamin C of 4T1 cells with doxycycline induced ATM depletion by shRNA (shATM) and non-target (shNT) controls following 4 days of treatment with doxycycline. Histone H3 was used as loading control. **(B)** Corresponding quantification of the lamin A and C levels based on three independent experiments. Lamin levels were normalized to shNT cells. *, *p* < 0.05 based on Two-way ANOVA with Tukey’s multiple comparison test. *N* = 3. **(C)** Representative immunoblot for lamin A and lamin C of MEFs with *Atm* deletion (*Atm*-Null) and wild-type (WT) controls. Histone H3 was used as loading control. **(D)** Corresponding quantification of lamin A and C protein levels from immunoblots based on three independent experiments. Lamin levels were normalized to WT controls. ****, *p* < 0.0001 based on two-way ANOVA with Tukey’s multiple comparison test. *N* = 3. **(E)** Representative image panel of immunofluorescence staining for lamin A/C of *Atm*-Null and wild-type (WT) MEFs. Scale bar: 20 µm **(F)** Corresponding quantification of the lamin A/C total immunofluorescence intensity per nucleus, based on the mean values from three independent experiments. *N* = 3; representing a total of 184 cells for wild-type and 154 cells for *Atm*-Null. ****, *p* < 0.0001, based on unpaired *t*-test with Welch’s correction. **(G)** Quantification of nuclear size of *Atm*-Null and WT MEFs. **, *p* < 0.01, based on the mean values of three independent experiments, using unpaired *t*-test with Welch’s correction. Error bars in this figure represent mean ± s.e.m. of the mean values of each experiment.

### ATM regulates lamin levels transcriptionally and not post-translationally

To evaluate the possible mechanism by which ATM regulates lamin A protein levels, we investigated the impact of ATM on *LMNA* gene expression. Quantitative real-time PCR analysis revealed that *Atm*-Null MEFs had significantly lower *LMNA* mRNA expression compared to wild-type controls (Fig. 3A). Similar results were obtained in 4T1 mouse mammary carcinoma cells following inducible ATM depletion (Fig. 3B). Thus, our findings suggest that deletion or depletion of ATM reduces *LMNA* gene expression, thereby decreasing lamin A/C protein levels. Since the primers used for the PCR analysis target a region of the *LMNA* gene common to both lamin A and C, we could not distinguish whether transcripts for lamin A and C were differentially regulated.

**Figure 3:**
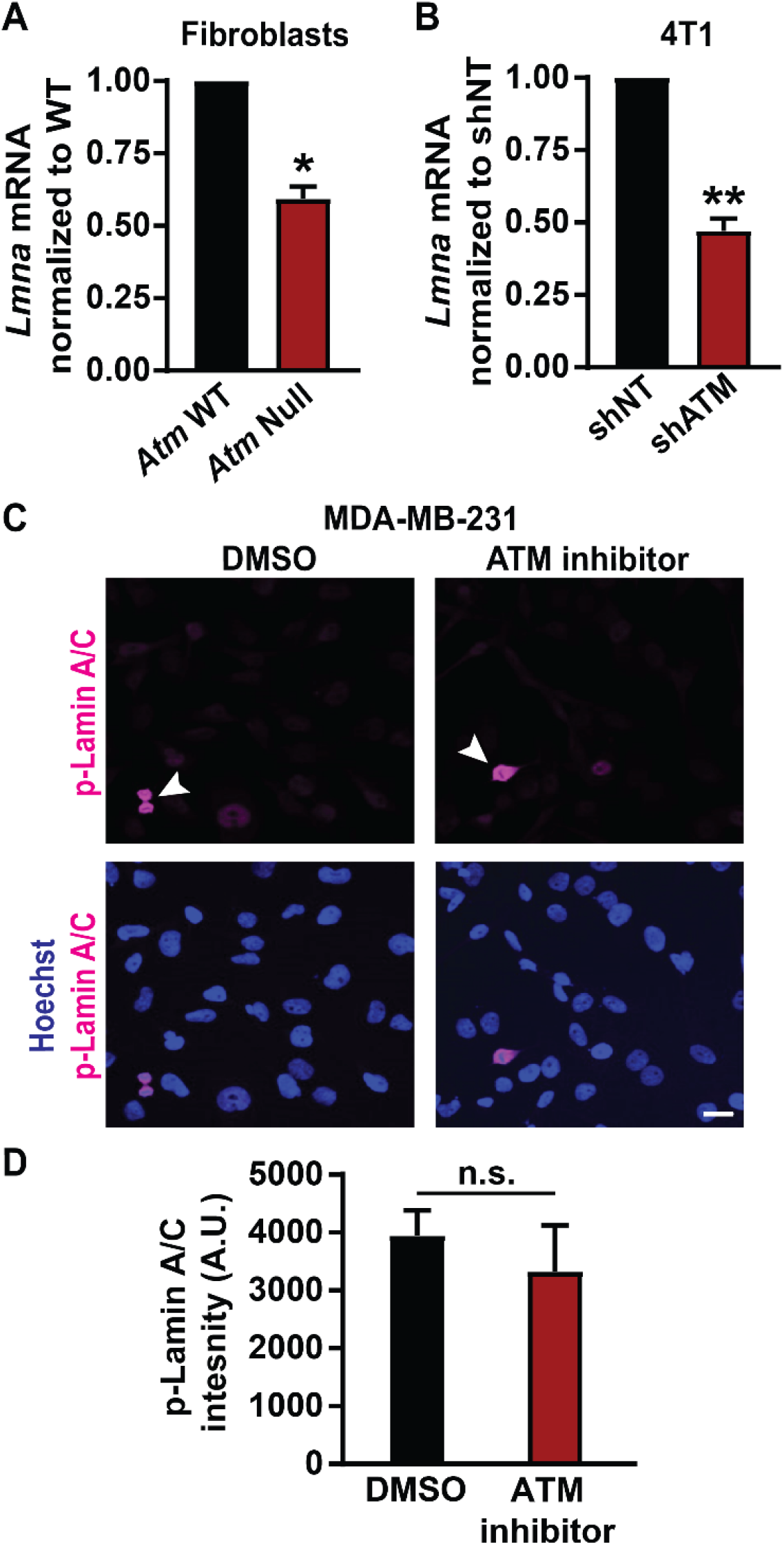
Loss of ATM results in decreased lamin A/C transcription. **(A)** *LMNA* gene transcription levels in *Atm* wild-type (WT) and *Atm*-Null MEFs quantified by Real-time PCR, based on three independent experiments. Results were normalized to *Atm* WT cells. *, *p* < 0.05 based on unpaired *t*-test with Welch’s correction. *N* = 3. **(B)** *Lmna* gene transcription levels in 4T1 cells with inducible depletion of *Atm* by shRNA (shATM) and non-target controls (shNT). Results were normalized to shNT cells. **, *p* < 0.01 based on unpaired *t*-test with Welch’s correction. *N* = 3 **(C)** Representative image panel of immunofluorescence labeling for phospho-lamin A/C (Ser22) in MDA-MB-231 cells treated with 0.1% DMSO or 10 µM ATM inhibitor KU-55933. Mitotic cells (arrowhead) within the cell population serve as positive controls. Scale bar: 20 µm. **(D)** Corresponding quantification of phospho-lamin A/C total immunofluorescence intensity, based on the average values from three independent experiments. *N* = 3; unpaired *t*-test with Welch’s correction. Error bars in this figure represent mean ± s.e.m. of the mean values of each experiment.

To determine whether post-translational modifications of lamins could additionally contribute to reduced lamin protein levels, we examined whether ATM inhibition increases phosphorylation of lamin A/C. Phosphorylation of lamin A/C at residues Ser22 and Ser329 by Cdk1 promotes nuclear lamina disassembly and degradation (Gerace and Blobel, 1980; Heald and McKeon, 1990; Peter et al., 1990; Swift et al., 2013; Buxboim et al., 2014) and could explain the observed reduction in lamin A protein levels. However, MDA-MB-231 cells treated with the ATM inhibitor KU-55933 had similar levels of phospho-lamin A/C (pSer22) as vehicle-treated control cells (Fig. 3C, D). Except for mitotic cells (Fig. 3C, arrowheads), which showed the expected increase in phosphorylated lamin A/C (Heald and McKeon 1990) and thus served as positive controls for the pSer22 antibody, both the ATM inhibitor treated and the vehicle control treated cells had very low pSer22 levels (Fig. 3C) that were not statistically different (Fig. 3D). These findings suggest that reduced lamin levels due to ATM inhibition are not due to increased phosphorylation of lamin A/C. Taken together, our data suggest that ATM modulates lamin A levels transcriptionally and not through increased phosphorylation, although we cannot exclude that ATM inhibition or deletion could alter the stability or translation of *LMNA* transcripts, for example via miRNAs, or promote lamin A protein degradation by altering other post-translational modifications.

### ATM depletion or inhibition increase nuclear deformability

Lamin A/C is a key contributor for nuclear structure and stability (Broers et al., 2004; Lammerding et al., 2004; Bell and Lammerding, 2016; Dahl and Aird, 2017). Since inhibition or depletion of ATM reduces lamin A/C levels, we hypothesized that these ATM manipulations may also result in more deformable nuclei. To measure nuclear deformability, we used a custom-built microfluidic micropipette aspiration device, wherein the length of nuclear protrusions of cells inside 3 × 5 µm^2^ aspiration channels subjected to a precisely controlled externally applied pressure (Fig. 4A) can be directly correlated to nuclear deformability (Davidson et al., 2019). Micropipette aspiration analysis revealed that MDA-MB-231 cells treated with an ATM inhibitor had higher nuclear protrusion lengths compared to vehicle-treated control cells (Fig. 4A-B), indicating that ATM inhibition indeed increases nuclear deformability. To confirm the effect using genetic deletion of *Atm*, we compared *Atm*-Null with wild-type controls MEFs in the microfluidic micropipette aspiration assay (Fig. 4C; Suppl. Video 1). *Atm*-Null MEFs had significantly increased nuclear protrusions inside the micropipette channels compared to wild-type control cells (Fig. 4C-D), indicative of more deformable nuclei, similar to the effect observed during ATM inhibition in human cells. Taken together, our findings indicate that chemical inhibition or genetic deletion of ATM leads to more deformable nuclei due to the reduced lamin A levels in these cells.

**Figure 4:**
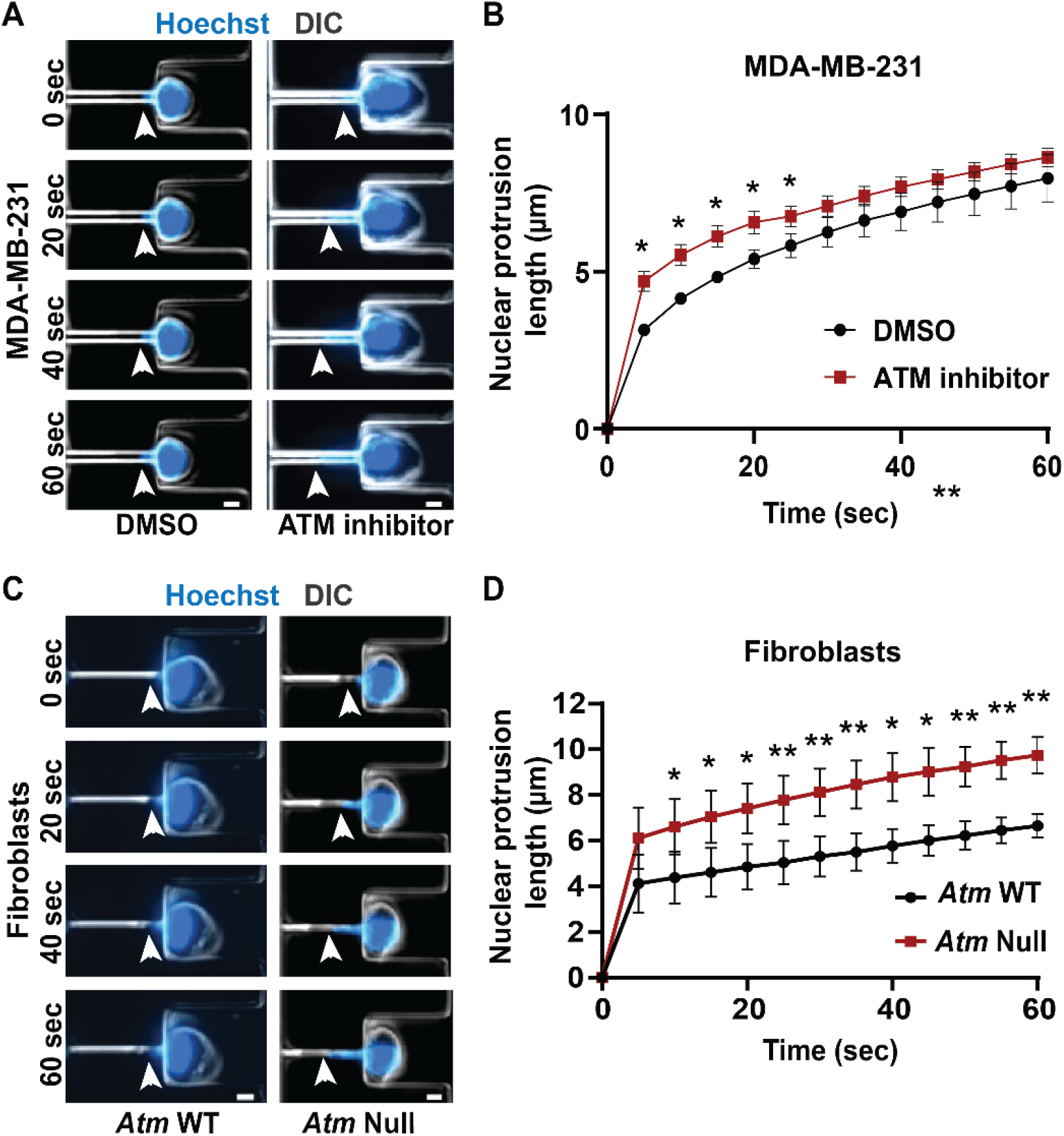
Functional loss of ATM makes nuclei more deformable. **(A)** Representative time-lapse image sequence of MDA-MB-231 cells treated with ATM inhibitor KU-55933 (*n* = 95 cells) or vehicle control (DMSO, *n* = 45 cells) undergoing nuclear deformation in micropipette aspiration device with channels of 3 × 5 µm^2^ in size. Arrowheads indicate the leading edge of nuclear protrusion. Scale bar: 5 µm **(B)** Quantification of the nuclear protrusion length inside the micropipette aspiration channels as shown in (A) based on the means of three independent experiments. *, *p* < 0.05; based on paired *t*-test, *N* = 3. **(C)** Representative image panel of *Atm*-Null MEFs (*n* = 117 cells total) and *Atm* wild-type (WT) controls (*n* = 130 cells total) undergoing nuclear deformation in micropipette aspiration device with channels of 3 × 5 µm^2^ in size. Arrowheads indicate the leading edge of nuclear protrusion. Scale bar: 5 µm **(D)** Quantification of the nuclear protrusion length inside the micropipette aspiration channels as shown in (C), based on the means of three independent experiments. *, *p* < 0.05; **, *p* < 0.01; based on paired *t*-test, *N* = 3. Error bars in this figure represent mean ± s.e.m. of the mean values of each experiment.

### Cells lacking ATM migrate faster through confined spaces

Nuclear deformability is a key factor regulating migration through confined spaces (Wolf et al., 2013; Davidson et al., 2014; Harada et al., 2014; Lautscham et al., 2015; Denais et al., 2016). Cancer cells, immune cells, stem cells, and fibroblasts regularly squeeze through tight interstitial spaces of the order of 1-20 µm in diameter when they migrate through the extracellular matrix and tissues (Doerschuk et al., 1993; Stoitzner et al., 2002; Weigelin et al., 2012). Successful migration through such confined spaces is largely dependent on the ability of the nucleus to deform through the available space (Balzer et al., 2012; Fu et al., 2012; Tong et al., 2012; Wolf et al., 2013; Davidson et al., 2014; Lautscham et al., 2015) and lack of lamin A/C allows faster confined migration (Davidson et al., 2014; Harada et al., 2014).Since depletion or inhibition of ATM reduces lamin A levels and makes nuclei more deformable, we hypothesized that depletion or inhibition of ATM may result in faster cell migration through confined spaces.

To evaluate confined migration efficiency, we performed time-lapse microscopy of cells migrating through custom-developed microfluidic devices that mimic the confined spaces found in tissues (Davidson et al., 2015; Keys et al., 2018). These devices contain channels with small constrictions (either 1 × 5 µm^2^ or 2 × 5 µm^2^ in cross-section) that require extensive nuclear deformation, and larger (15 × 5 µm^2^) control channels, in which the nucleus does not have to deform substantially (Davidson et al., 2014; Davidson et al., 2015; Denais et al., 2016; Elacqua et al., 2018). To monitor cells as they migrate through the microfluidic devices (Fig.5A-B; Suppl. Videos 2 & 3) and determine their transit time through the constrictions or the larger control channels, cells were modified to stably express a green fluorescent protein with a nuclear localization sequence (NLS-GFP) (Denais et al., 2016; Elacqua et al., 2018). MDA-MB-231 and HT1080 cells treated with an ATM inhibitor migrated faster through the small constrictions than vehicle-treated control cells (Fig. 5A-B, E-F). In contrast, ATM inhibition did not alter migration through the larger control channels that do not require large nuclear deformation (Fig. 5C-D), suggesting that ATM inhibition affects specifically migration through confined spaces and not overall migration efficiency. This effect can likely be attributed to the reduced lamin A levels and increased nuclear deformability following ATM inhibition. We corroborated these results in experiments comparing *Atm*-Null MEFs with wild-type controls migrating through the microfluidic devices, which showed that *Atm*-Null MEFs exhibited shorter transit times through the small constrictions compared to wild-type control MEFs but did not exhibit any significant differences in migration through the larger control channels (Fig. 5G). Intriguingly, the Atm-Null MEFs showed a trend towards increased nuclear envelope rupture inside the small constrictions (Fig. 5H), consistent with previous findings in lamin A/C-deficient cells (Denais et al., 2016; Raab et al., 2016), although this trend did not reach statistical significance (*p* = 0.11). Collectively, these results indicate that deletion or inhibition of ATM increases cell migration speed through confined spaces by increasing nuclear deformability.

**Figure 5:**
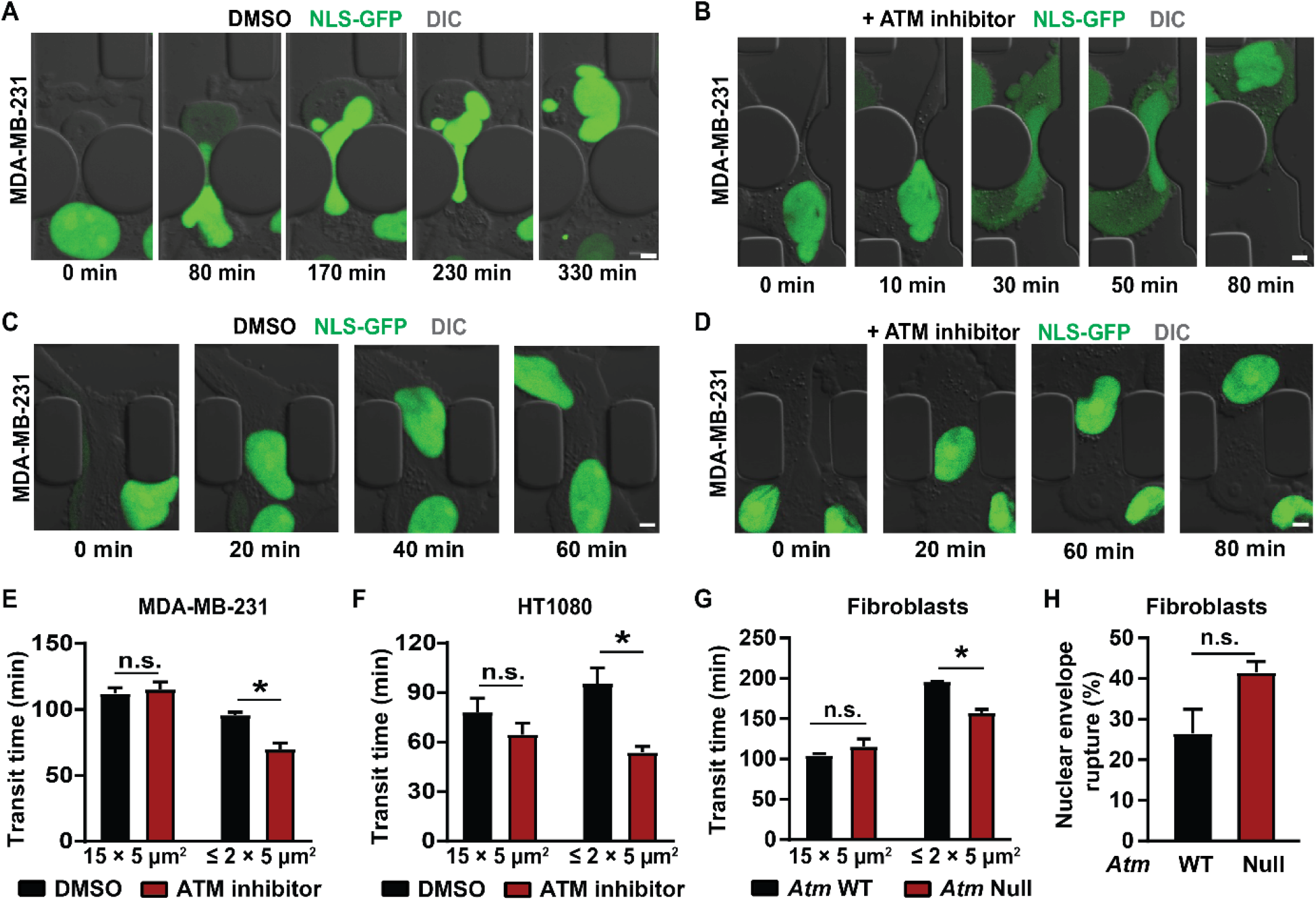
Cells lacking ATM migrate faster through confined spaces. **(A)** Representative time-lapse image sequences of an MDA-MB-231 breast cancer cell expressing NLS-GFP and treated with vehicle control (DMSO) during migration through a 1 × 5 µm^2^ constriction in the microfluidic device. Scale bar: 5 µm. **(B)** Representative image sequence of an MDA-MB-231 breast cancer cell expressing NLS-GFP and treated with an ATM inhibitor (KU-55933) during migration through a 1 × 5 µm^2^ constriction in the microfluidic device. Scale bar: 5 µm. **(C)** Representative time-lapse image sequences of an MDA-MB-231 breast cancer cell expressing NLS-GFP and treated with vehicle control (DMSO) during migration through a 15 × 5 µm^2^ control channel in the microfluidic device. Scale bar: 5 µm. **(D)** Representative image sequence of an MDA-MB-231 breast cancer cell expressing NLS-GFP and treated with an ATM inhibitor (KU-55933) during migration through a 15 × 5 µm^2^ control channel in the microfluidic device. Scale bar: 5 µm. **(E)** Quantification of transit times for MDA-MB-231 cells to migrate through ≤ 2 × 5 µm^2^ constrictions or 15 × 5 µm^2^ control channels when treated with ATM inhibitor KU-55933 or vehicle control (DMSO). *, *p* < 0.05 based on two-way ANOVA with Tukey’s multiple comparison test; *N* = 3 for each group, representing the means of three independent experiments; total number of cells analyzed: 114 cells for DMSO and 83 cells for ATM inhibitor for the small constrictions; 72 cells for DMSO and 38 cells for ATM inhibitor for the larger control channels. **(F)** Transit times for HT1080 fibrosarcoma cells to cross the ≤ 2 × 5 µm^2^ constrictions or 15 × 5 µm^2^ control channels when treated with ATM inhibitor KU-55933 or vehicle control (DMSO). *, *p* < 0.05 based on Two-way ANOVA with Tukey’s multiple comparison test; *N* = 3 for each group, representing the means of three independent experiments; total number of cells analyzed: 200 cells for DMSO and 276 cells for ATM inhibitor for the small constrictions; 182 cells for DMSO and 140 cells for ATM inhibitor for the larger control channels. **(G)** Transit times for *Atm* wild-type (WT) or *Atm*-Null MEFs to cross the ≤ 2 × 5 µm^2^ constrictions or the 15 × 5 µm^2^ control channels. *, *p* < 0.05 based on Two-way ANOVA with Tukey’s multiple comparison test; *N* = 3 for each group, representing the means of three independent experiments; total number of cells analyzed: 336 *Atm* WT and 221 *Atm*-Null MEFs for the small constrictions; 133 *Atm* WT and 111 *Atm*-Null MEFs for the larger control channels. **(H)** Quantification of nuclear envelope rupture events of Atm WT and Atm-Null MEFs migrating through the ≤ 2 × 5 µm^2^ constrictions. The difference was not statically significant (*p* = 0.11, based on the means of three independent experiments. *N* = 3. Error bars in this figure represent mean ± s.e.m. of the mean values of each experiment.

To test whether other DDR kinases can also affect migration through confined spaces, we treated MDA-MB-231 and HT1080 cells with inhibitors for either ATR or DNA-PK and assessed migration efficiency in the microfluidic devices. Inhibition of ATR or DNA-PK did not significantly increase migration speed through the constrictions in either MDA-MB-231 or HT1080 cells (Suppl. Fig. 3). These findings indicate that only ATM inhibition or deletion promotes confined migration, consistent with our finding that neither ATR nor DNA-PK significantly altered lamin A levels (Fig. 1D-I). Collectively, our studies indicate a novel role for ATM in modulating nuclear lamin A levels, wherein deletion or inhibition of ATM leads to reduced lamin A levels by reducing *LMNA* transcription, thereby resulting in increased nuclear deformability and increased migration speed through confined spaces.

## Discussion

Using a comprehensive approach with chemical inhibitors, stable and conditional genetic depletion of ATM across multiple cell lines, we showed that inhibition or depletion of ATM, but not ATR or DNA-PK, leads to reduced lamin A levels and increased nuclear deformability. This effect is due to reduced *LMNA* gene expression in cells depleted for ATM, although we cannot rule out that additional mechanisms may contribute to the reduced lamin A levels, including miRNAs and/or posttranslational modifications that lead to increased lamin A/C degradation. We have thus uncovered a novel role for ATM outside of its activities in DNA repair, in regulating nuclear stiffness and mechanics by modulating nuclear lamin levels. In addition, we have demonstrated that the increased nuclear deformability resulting from ATM inhibition or deletion enables cells to migrate faster through confined spaces without affecting migration speed through larger spaces that do not require substantial nuclear deformation, suggesting that lack of ATM specifically affects confined migration but not migration in general.

Interestingly, whereas our study uncovered a new role for ATM in modulating lamin A levels, we did not see a similar impact on lamin expression and confined migration upon inhibiting other DDR kinases, such as ATR and DNA-PK (Figs. 1, 5; Suppl. Fig. 3). This finding suggests that regulation of lamin A levels is likely independent of ATM’s DNA repair activities, where ATM shares structural and functional similarities with ATR and DNA-PK (Perry and Kleckner, 2003; Lempiainen and Halazonetis, 2009; Lovejoy and Cortez, 2009; Awasthi et al., 2015; Blackford and Jackson, 2017). This interpretation is supported further by a previous study, which found that although ATR mutations and depletion lead to altered nuclear morphology and stability, these defects were not mediated by changes in lamin A/C expression or localization (Kidiyoor et al., 2020). Although we show that ATM reduces lamin A levels transcriptionally and not through increased phosphorylation (Fig. 3), the exact mechanism by which ATM modulates lamin A levels remains to be determined, including the more pronounced reduction of lamin A compared to lamin C in the human cell lines (Fig. 1A-C), and the surprising finding that lamin A/C depletion was associated with increased lamin B1 levels in the *Atm*-Null MEFs (Suppl. Fig. 1C).

One possible pathway by which ATM could affect lamin A/C expression is through ATM’s interaction with the Akt pathway (Halaby et al., 2008; Li and Yang, 2010). Akt is a major signalling hub involved in the regulation of cellular metabolism, survival, growth as well as cell cycle progression (Manning and Cantley, 2007). ATM has previously been reported to activate Akt in response to DNA damage or insulin treatment (Viniegra et al., 2005; Halaby et al., 2008; Li and Yang, 2010; Liu et al., 2014). Although Akt has numerous targets, previous studies have shown that Akt can specifically regulate *LMNA* gene expression and degradation (Bertacchini et al., 2013; Naeem et al., 2015). Thus, it is possible that deletion of ATM reduces Akt activity and thereby downregulates *LMNA* gene transcription, resulting in lower lamin A/C levels. Notably, lamin A/C is a substrate for Akt, and Akt mediated phosphorylation of lamin A/C can promote lamin A/C for degradation (Cenni et al., 2008; Bertacchini et al., 2013; Naeem et al., 2015). However, in our experiments, we did not observe any significant difference in lamin A/C phosphorylation following ATM inhibition (Fig. 3C-D). Interpreting a potential role of Akt in modulating lamin A/C levels is further complicated by the fact that Akt activity can lead to both increases and decreases in lamin A/C levels, depending on the specific context. A previous study found that overexpression of Akt resulted in a more than 4-fold increase of lamin A in interphase cells, but a 50% decrease in lamin A expression in mitotic cells (Bertacchini et al., 2013). In recent experiments with MDA-MB-231 cells, we found that Akt inhibition resulted in a significant increase in lamin A/C protein levels, without changing *LMNA* gene expression (Bell et al., 2021). Thus, it remains to be investigated if and how Akt may mediate effects of ATM inhibition on lamin A/C protein levels.

ATM could also modulate lamin A/C transcription through additional pathways. ATM can regulate epigenetic modifications via its interactions with histone deacetylases (HDACs) (Kim et al., 1999). HDACs promote heterochromatin formation and have been shown previously to regulate gene expression in an ATM dependent manner (Jang et al., 2010) and to interact with lamin A/C (Mattioli et al., 2018; Mattioli et al., 2019). Moreover, ATM regulates KAP1 (KRAB domain associated protein), a known transcriptional co-repressor (White et al., 2006; Callen et al., 2009) and promotes histone H2A ubiquitination, a histone modification linked to transcription repression in response to DNA damage (Shanbhag et al., 2010). Thus, interaction of ATM with HDACs, H2A ubiquitylation, or KAP1 could constitute other possible mechanisms by which ATM regulates *LMNA* gene transcription that should be investigated in future studies. Whether these effects are specific to *LMNA* expression or could also affect other nuclear envelope proteins remains to be determined. Future studies should be directed to assess the transcriptomic consequences of ATM inhibition or deletion, which may identify key pathways or regulators mediating lamin A/C expression.

### Limitations of current study

As discussed above, one of the major limitations of this study is that the precise mechanism by which deletion or inhibition of ATM causes reduced levels of lamin A remains to be elucidated. Confounding this, although we found that deletion or depletion of ATM reduced levels of both lamin A and C in MEFs and mouse mammary carcinoma cells, inhibition of ATM in human cancer cells substantially reduced lamin A levels, whereas the reduction in lamin C levels was not statistically significant. It remains unclear whether this difference is due to the different approaches to manipulate ATM levels/function or whether it represents species-specific differences, as both 4T1 and MEFs are murine cells, whereas MDA-MB-231 and HT1080 cells are human. Notably, we were unable to distinguish the impact of ATM depletion on lamin A and lamin C separately in some of the assays (like the immunofluorescence staining and Real-Time PCR) as these proteins are splice variants of the same *LMNA* gene and thus share a large common sequence identity, except for a short unique C-terminal section. Furthermore, for *Atm*-Null MEFs, where *Atm* deletion modulated both lamin A and lamin C levels (Fig. 2 C-D) we could not separate the effect of lamin A and lamin C reduction independently on nuclear deformation and migration speed in our assays. Moreover, the antibody used to detect B-type lamins was specific to lamin B1, preventing us from making any inferences on the effect of lamin B2 levels.

It will also be important to further elucidate the effect of ATM inhibition or depletion on specific functions mediated by lamin A. We demonstrated that ATM inhibition and deletion increase nuclear deformability and enhance migration through confined spaces. However, in addition to modulating nuclear stiffness and stability, lamins A and C also have diverse other functions, including in gene regulation, cytoskeletal organization, proliferation, and differentiation (Ho and Lammerding, 2012; Schreiber and Kennedy, 2013; Davidson et al., 2014; Bell and Lammerding, 2016). The impact of ATM deletion or inhibition on these lamin functions remains to be determined. It is also intriguing to speculate about the effect of specific ATM mutations, including those responsible for A-T, on lamin levels. Unfortunately, we were unable to obtain patient-derived cells to directly assess this effect.

Of note, ATM and lamins independently affect major metabolic and DNA repair pathways (Singh et al., 2013; Gonzalo, 2014; Awasthi et al., 2015; Blackford and Jackson, 2017; Charar and Gruenbaum, 2017; Dahl and Aird, 2017) and future studies on understanding the impact of this ATM – lamin A crosstalk on these pathways will be needed. Studies done in mouse models of lamin A/C-related progerias reveal extensive connections between lamin A/C and ATM. For example, progeroid *Lmna* mutations cause ATM-dependent activation of NF-κB signaling and pro-inflammatory cytokine secretion (Osorio et al., 2012), and ATM inhibition in a progeria cell model resulted in increased mitochondrial function, reduced senescence, reduced accumulation of mutant lamin A, and improved nuclear shape (Kuk et al., 2019).

Lastly, while our present investigation focused on the impact of ATM inhibition and depletion on lamin A/C levels, the impact of lamin A levels on ATM expression and function remains unknown. Recent studies found that lamin A/C depletion results in impaired DNA base excision repair and reduced expression of several enzymes mediating base excision repair (Maynard et al., 2019), and that lamin A/C depletion impaired the recruitment of proteins required to protect single strand DNA at stalled replication forks (Graziano et al., 2021). However, these studies did not investigate the effect of lamin A/C depletion on ATM function. Given the emerging cross-talk between lamins and DDR pathways (Graziano et al., 2018; Sengupta et al., 2020), future studies should address whether ATM and lamins positively cross-regulate each other.

### Conclusions

Despite these limitations, our findings identify a new link between ATM and lamins that could potentially explain the impact these proteins have on different cellular pathways including metabolism, oxidative stress, and DNA repair. Insights from this work will be relevant to cells carrying mutations in the *ATM* gene, which are present in many cancers (e.g., in about 45% of lymphomas)(Choi et al., 2016), wherein lack of ATM may promote invasion or intra- and extravasation of tumor cells during the early steps of metastasis (Chaffer and Weinberg, 2011; Davidson et al., 2014). Furthermore, our findings may have wide implications for patients suffering from A-T. Mutations and deletions in the *LMNA* gene have been shown previously to promote oxidative stress, premature ageing and neurological disorders (Rankin and Ellard, 2006). Thus, reduced lamin A levels due to non-functional ATM could explain the neurological disorders and premature ageing characteristic of this disease (Boder and Sedgwick, 1958). Collectively, we have identified a new role of ATM in modulating lamin A levels, thereby altering nuclear stiffness and cell migration through confined spaces, which has broad biological and biomedical implications and thus the potential to spawn numerous further investigations.

## Materials and Methods

### Cells and cell culture

The breast adenocarcinoma cell line MDA-MB-231 (ATCC HTB-26) was purchased from American Type Culture Collection (ATCC); the fibrosarcoma cell line HT1080 (ACC 315) was a gift from Peter Friedl and Katarina Wolf and originally purchased from the DSMZ Braunschweig, Germany; the *Atm* wild-type (WT) and *Atm*-Null mouse embryonic fibroblasts (MEFs) were a gift from Robert Weiss (Balmus et al., 2012). The BALB/c-derived mammary carcinoma cell line 4T1 was obtained from Dr. Miller at Karmanos (Aslakson and Miller, 1992) in 2002, and authenticated by IDEXX Bioresearch (Columbia, MO, USA) in 2019. 4T1shNT and 4T1shATM were prepared as described below. All cells were cultured in Dulbecco’s Modified Eagle Medium (DMEM) supplemented with 10% (v/v) fetal bovine serum (FBS, Seradigm VWR) and 1% (v/v) penicillin and streptomycin (PenStrep, ThermoFisher Scientific) and under humidified conditions at 37°C and 5% CO_2_. 4T1shNT and 4T1shATM cells were cultured with 4 µg/ml doxycycline for 4 days to induce shRNA mediated ATM depletion, prior to each experiment. The human cancer cells and the *Atm* MEFs were stably modified with lentiviral vectors to express the nuclear rupture reporter NLS-GFP (pCDH-CMV-NLS-copGFP-EF1-blastiS, available through Addgene (#132772)) (Denais et al., 2016). The identity of human cell lines MDA-MB-231 and HT1080 was confirmed using ATCC’s STR profiling service.

### Drug treatment

For experiments with DDR inhibitors, MDA-MB-231 and HT1080 cells were treated with either DMSO (final concentration of 0.1% by volume) or ATM inhibitor (KU55933, final concentration of 10 µM), ATR inhibitor (VE-821, final concentration of 10 µM) or DNA-PK inhibitor (NU-7441, final concentration of 5 µM) dissolved in DMSO, starting 48 hours prior to collecting cell lysates.

### Viral modification

For the production of cells stably expressing NLS-GFP, pseudoviral particles were produced as described previously (Hanson et al., 2008). In brief, 293-TN cells (System Biosciences, SBI) were co-transfected with the lentiviral plasmid and lentiviral helper plasmids (psPAX2 and pMD2.G, gifts from Didier Trono) using PureFection (SBI), following manufactures protocol. Lentivirus containing supernatants were collected at 48 hours and 72 hours after transfection and filtered through a 0.45 µm filter. Cells were seeded into 6-well plates so that they reached 50-60% confluency on the day of infection and transduced at most 3 consecutive days with the viral stock in the presence of 8 µg/mL polybrene (Sigma-Aldrich). The viral solution was replaced with fresh culture medium, and cells were cultured for 24 hours before selection with 6 μg/mL of blasticidine S (InvivoGen) for 10 days. Cells were sorted using fluorescence assisted cell sorting (FACS) to ensure expression of all the fluorescent reporters and maintained in media containing antibiotics (6 µg/mL of Blasticidin S) to ensure continued plasmid expression. For the production of 4T1 cells with stable expression of shRNA targeting *Atm* or a non-target control, HEK 293-FT cells were co-transfected with packaging plasmids pPAX2 and pMD2 and a pTRIPZ vector containing a tetracycline-inducible promoter (GE Dharmacon technology, kindly provided by Dr. Robert Schneider) driving the shRNA expression. shRNA directed against *Atm* mRNA (mouse-shRNA: ACTACATCAACTGCTTACTAAG) or a non-targeting sequence (NT; mouse-human-shRNA: AATTCTCCGAACGTGTCACGT) were cloned into pTRIPZ using EcoRI and XhoI restriction sites. 4T1 cells were transduced with cell-free virus-containing supernatants for 48 hours and selected with 100 µg/mL of neomycin (Gibco) for 96 hours.

### Fabrication and use of microfluidic migration devices

The devices were prepared as described previously (Davidson et al., 2014; Davidson et al., 2015). Microfluidic devices were first assembled by plasma treating the PDMS pieces and glass coverslips (pretreated with 0.2 M HCl) for 5 min, then immediately placing the PDMS pieces on the activated coverslips and gently pressing to form a covalent bond. The finished devices were briefly heated on a hot plate at 95°C to improve adhesion. Devices were filled with 70% ethanol, then rinsed with autoclaved deionized water and coated with extracellular matrix proteins. For all cell lines, except MEFs, devices were coated with 50 µg/mL type-I rat tail collagen (BD Biosciences) in acetic acid (0.02 N) overnight at 4°C. For MEFs, devices were incubated with fibronectin (Millipore) in PBS (2 to 20 µg/mL) overnight at 4°C. After the incubation, devices were rinsed with PBS and cell culture medium before loading the cells into the devices (about 50,000-80,000 cells per chamber). Subsequently, devices were placed inside a tissue culture incubator for a minimum of 24 hours to allow cell attachment before mounting the devices on a microscope for live-cell imaging. The media reservoirs of the device were covered with glass coverslips to minimize evaporation during live cell imaging. Cells were imaged every 5-10 min for 14-16 hours in phenol-red free medium FluoroBrite DMEM supplemented with 10% FBS.

For MEFs, platelet derived growth factor (PDGF) was used as a chemoattractant at a concentration of 200 ng/ml in the well towards which cells were migrating. Transit times were quantified by manual analysis. Briefly, transit time of a cell through a constriction was defined as the time between when the front of the cell crossed an imaginary line 5 µm in front of the constriction center and when the back of the nucleus passed a corresponding line 5 µm behind the constriction center. A similar analysis was performed on cells in the 15 µm wide channels to obtain values of the migration transit time of cells under non-constrained conditions. For experiments with DDR inhibitors, MDA-MB-231 and HT1080 cells were treated with either DMSO alone (Sigma, final concentration of 0.1% by volume) or ATM inhibitor (KU55933, final concentration of 10 µM), ATR inhibitor (VE-821, final concentration of 10 µM), or DNA-PK inhibitor (NU-7441, final concentration of 5 µM) dissolved in DMSO starting 24 hours prior to the start of migration experiments. The treatment was continued for the length of the experiment. Transit times were calculated using a custom-written MATLAB program as described previously (Elacqua et al., 2018).

### Fabrication and use of micropipette aspiration devices

The devices were prepared as described previously (Davidson et al., 2019). In brief, PDMS molds of the devices were cast using Sylgard 184 (Dow Corning). Three ports were made using a 1.2-mm biopsy punch and the device was bonded to glass coverslips using plasma treatment. Pressure at the inlet and outlet ports was set to 1.0 and 0.2 pounds per square inch (relative to atmospheric pressure, *P*_atm_), respectively, using compressed air regulated by a MCFS-EZ pressure controller (Fluigent) to drive single cells through the device. The exit port was set to *P*_atm_ and outfitted with a hand-held pipette to flush cells from the pockets at the start of each image acquisition sequence. Cells (circa 5 × 10^6^ cells/ml) were suspended in 2% bovine serum albumin (BSA), 0.2% FBS and 10 μg/ml Hoechst 33342 (Invitrogen) DNA stain in PBS and were captured within an array of 18 pockets and then forced to deform into micropipette channels with 3 µm × 5 µm cross sections. Bright-field and fluorescence images were acquired every 5 seconds for 120 to 300 seconds. Nuclear protrusion length was calculated using a custom-written MATLAB program (Davidson et al., 2019), and used as read-out for nuclear deformability. For micropipette aspiration experiments with DDR inhibitors, MDA-MB-231 cells were pre-treated with either DMSO (final concentration of 0.1% by volume) or ATM inhibitor (KU55933, final concentration of 10 µM) dissolved in DMSO for 48 hours prior to the start of experiments.

### Western Blot analysis

For quantification of lamin A/C levels, cells were lysed in high salt RIPA buffer containing protease (complete EDTA-Free, Roche) and phosphatase (PhosSTOP, Roche) inhibitors. Protein was quantified using Bio-Rad Protein Assay Dye, and 20–30 µg of protein lysate was separated using a 4–12% Bis-Tris polyacrylamide gel with a standard SDS–PAGE protocol. Protein was transferred to a polyvinylidene fluoride membrane for 1 hour at room temperature at 16 V voltage. Membranes were blocked using 3% BSA in TBST, and primary antibodies (Lamin A/C (Santa Cruz) - dilution: 1:1000; lamin B1 (Abcam) – dilution 1:1000; H3 (Abcam) - dilution: 1:5000) were diluted in the same blocking solution and incubated overnight at 4 °C. Protein bands were detected using either IRDye 680LT or IRDye 800CW (LI-COR) secondary antibodies, imaged on an Odyssey CLx imaging system (LI-COR) and analyzed in Image Studio Lite (LI-COR). To assess *Atm* expression, 4T1 shNT and shATM cells were lysed in RIPA buffer (Sigma) containing protease inhibitor cocktail (Millipore Sigma) and quantified using the Pierce™ BCA Protein Assay Kit (Thermo Fisher Scientific). 50 µg of protein lysate was separated using a 8% SDS-PAGE gel at 100 V voltage. Protein was transferred to a polyvinylidene fluoride membrane (Millipore Sigma) overnight at 4ºC and 50mA current. Membranes were blocked using 5% milk in Tris-buffered saline containing 0.1% Tween-20 (TBST) for 1 hour and primary antibodies (mouse ATM (Santa Cruz) – dilution: 1:1000; Actin (Cell Signaling Technology) – dilution: 1:1000) were diluted in the same blocking solution and incubated overnight at 4 °C. Protein bands were detected using either ECL anti-mouse or ECL anti-rabbit (Cytiva, dilution: 1:5000) secondary antibodies, visualized using chemiluminescence (Perkin Elmer) and imaged through Azure c600 imager (Azure Biosystems).

### qPCR analysis

Cells were lysed and RNA was collected using the QIAGEN RNeasy kit. The collected RNA was treated with DNase to lyse residual genomic DNA using the TURBO DNA-free kit (Thermo Fisher). cDNA was prepared from 750 ng of purified RNA using the iSCRIPT cDNA synthesis kit (Bio-Rad). qPCR was performed on the LightCycler 480 qPCR system (Roche) using the LightCycler 480 SYBR Green I Master mix (Roche). *LMNA* gene expression was probed using the primer sequences in Table 1; *RN7SK, GAPDH, 18S* and *TMX4* gene expression levels were used as housekeeping controls. The primers (Integrated DNA Technologies) used for each of these genes are listed in Table 1. qPCR analysis was performed using the *C*_*t*_ method and the settings listed in Table 2.

**Table 1.**
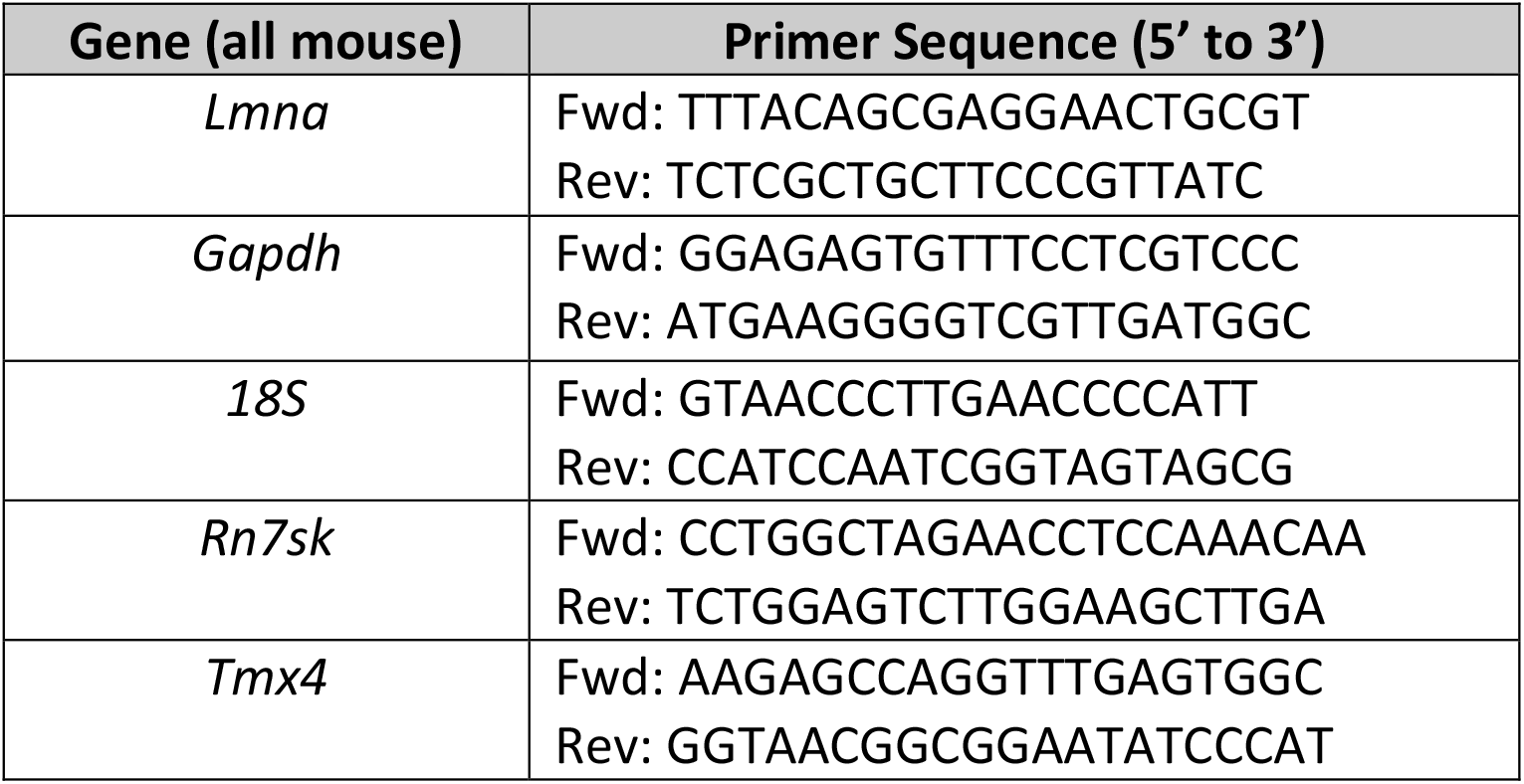
List of primers used for quantitative Real-Time PCR analysis of gene expression.

**Table 2.** Temperature and cycle settings for Real-Time PCR.

| Step | Cycles | Temperature (°C) | Duration |
| --- | --- | --- | --- |
| Pre-incubation | 1 | 95 | 10 min |
| Amplification | 50 | 95<br>60<br>72 | 15 sec<br>30 sec<br>30 sec |
| Melting curve | 1 | 95<br>50<br>97 | 5 sec<br>1 min<br>Continuous |
| Cooling | 1 | 40 | 30 sec |

### Immunofluorescence staining

Lamin A/C protein levels in the nucleus were quantified using immunofluorescence staining. Cells cultured on cover slips (pretreated with fibronectin (5 µg/ml in PBS) at 4°C overnight) were fixed with 4% PFA for 20 min at 37°C, permeabilized with PBS containing 0.25% Triton X-100 for 15 min at room temperature, washed, blocked with BSA and stained with anti-lamin A/C antibody (Santa Cruz Biotechnology (sc-376248), dilution 1:200) or anti-phopho lamin A/C antibody (Cell Signaling (#2026S), dilution 1:500) at 4°C overnight. Cells were washed with PBS and incubated for 1 h at room temperature with AlexaFluor antibodies (Invitrogen, dilution 1:300) and Hoechst 33342 (Invitrogen, dilution 1:1000) DNA stain. Fluorescence intensities were calculated based on total nuclear fluorescence, using a custom-written MATLAB program, made available on request. Nuclear size was quantified using ImageJ/FIJI (Schindelin et al., 2012).

### Fluorescence and confocal microscopy

Microfluidic device migration experiments and immunofluorescence staining were imaged on inverted Zeiss Observer Z1 microscope equipped with temperature-controlled stage (37°C) and CCD camera (Photometrics CoolSNAP KINO) using 20× air (NA = 0.8), 40× water (NA = 1.2) or 63× oil (NA = 1.4) immersion objectives. Airy units for all confocal images were set between 1.5 and 2.5. The image acquisition for migration experiments was automated through ZEN (Zeiss) software with imaging intervals for individual sections between 5-10 min. Migration experiments for MEFs were imaged on an IncuCyte (Sartorius) imaging system, using a 20× objective. Microfluidic micropipette experiments were collected on a motorized inverted Zeiss Observer Z1 microscope equipped with either charge-coupled device cameras (Photometrics CoolSNAP EZ or Photometrics CoolSNAP KINO) or a sCMOS camera (Hamamatsu Flash 4.0) using a 20× air (NA = 0.8) objective.

### Statistical analysis

All experimental results are based on at least three independent experiments. Unless otherwise noted, statistical analysis and error bars are based on the means of three independent experiments to account for variations from experiment to experiment. For data with normal distribution, we used either two-sided Student’s *t*-tests (when comparing two groups) or one-way analysis of variance (ANOVA) (for experiments with more than two groups) with post hoc tests. For experiments with two variables and more than two groups, two-way ANOVA was used with post hoc tests. All tests were performed using GraphPad Prism. Welch’s correction for unequal variances was used with all *t*-tests, comparing two groups while post hoc multiple-comparisons testing with Tukey’s correction was used with all two-way ANOVA analysis. Unless otherwise indicated, data are displayed as mean ± s.e.m., based on the means of individual experiments. For completeness, we have also listed the total number of cells included in the analysis. Data supporting the findings of this study are available from the corresponding author on reasonable request.

## Acknowledgements

The authors thank Darshil Safari, Catalina Pereira, Dongsun Kim, Jennie Sims for help with cells and reagents and members of the Lammerding lab for helpful discussion and support.The authors also thank Marcus Smolka for providing reagents and advice. This work was performed in part at the Cornell NanoScale Science & Technology Facility, a member of the National Nanotechnology Coordinated Infrastructure, which is supported by the National Science Foundation (Grant NNCI-2025233). This work was supported by awards from the National Institutes of Health (R01 HL082792, R01 GM137605, U54 CA210184 to J.L., R03 HD058220 to R.S.W., R01 CA201246 to S.D.) the Department of Defense Breast Cancer Research Program (Breakthrough Award BC150580 to J.L.), the National Science Foundation (URoL-2022048 to J.L.), the VolkswagenStiftung (A130142 to J.L.), and seed funding from the Cornell University Academic Integration program (to S.D., R.S.W., and J.L.). The content of this manuscript is solely the responsibility of the authors and does not necessarily represent the official views of the National Institutes of Health.

## Author Contributions

P.S. and J.L. conceptualized and designed the study. P.S., C.W.M., S.C. and C. V.-B. performed and analyzed all experiments. C. V.-B. and S.D. generated the *Atm* shRNA and shNT 4T1 cells. R.S.W contributed *Atm*-null and control MEFs plus crucial scientific guidance and advice on ATM and DDR pathways. P.S. and J.L. wrote the manuscript. All authors edited the manuscript.

## Potential conflicts of interest

SD has received compensation for consultant/advisory services from Lytix Biopharma, Mersana Therapeutics, EMD Serono, Ono Pharmaceutical, and Genentech, and research support from Lytix Biopharma and Boehringer-Ingelheim for unrelated projects. JL has provided consulting services for BridgeBio.

## Notes

### Summary of Updates

The manuscript has been revised to include additional data, replace the representative Western blots with higher quality images, and expanded the discussion section.

