## Supplementary Materials & Figures for "ATM modulates nuclear mechanics by regulating lamin A levels"

### Supplementary Figures

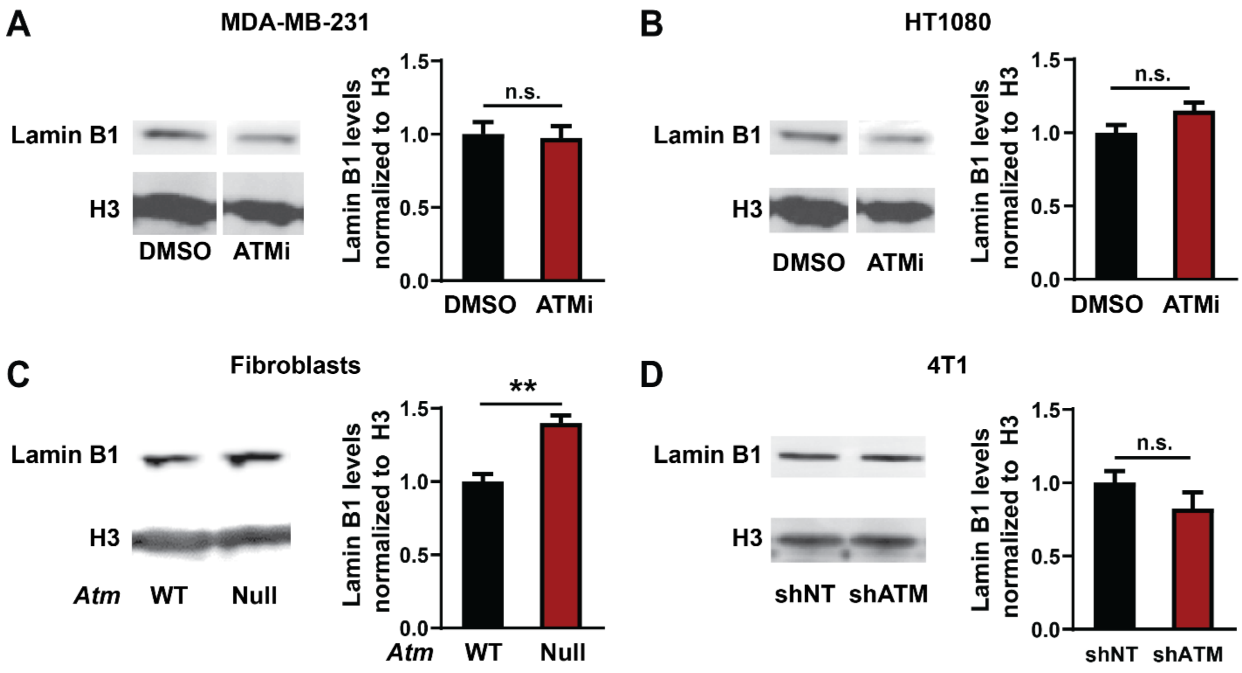


**Supplementary Figure 1: ATM inhibition or depletion does not alter lamin B1 levels. (A)** Representative immunoblot and quantification of lamin B1 protein levels in MDA-MB-231 cells treated with 10 µM of ATM inhibitor KU-55933 or 0.1% DMSO for 48 hours. Histone H3 was used as loading control. Lamin levels were normalized to DMSO treated cells. Differences are not statistically significant (n.s.), based on unpaired *t*-test with Welch's correction. $N = 3$. **(B)** Representative immunoblot and quantification for lamin B1 in HT1080 cells treated with 10 µM of ATM inhibitor KU-55933 or 0.1% DMSO for 48 hours. Histone H3 was used as loading control. Lamin levels were normalized to DMSO treated cells. Differences are not statistically significant (n.s.) based on unpaired *t*-test with Welch's correction. $N = 3$. **(C)** Representative immunoblot and quantification for lamin B1 in MEFs with *Atm* deletion (*Atm*-Null) and wild-type (WT) controls. Histone H3 was used as loading control. Lamin levels were normalized to WT controls. **, $p < 0.01$ based on unpaired *t*-test with Welch's correction. $N = 3$. **(D)** Representative immunoblot and quantification for lamin B1 in 4T1 cells with doxycycline inducible ATM depletion by shRNA (shATM) and non-target (shNT) controls. Histone H3 was used as loading control. Lamin levels were normalized to shNT cells. Differences are not statistically significant (n.s.) based on unpaired *t*-test with Welch's correction. $N = 3$. Error bars in this figure represent mean ± s.e.m.

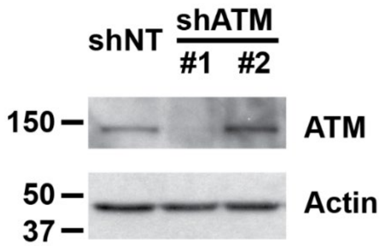


**Supplementary Figure 2: Depletion of ATM in 4T1 mouse mammary carcinoma cells using shRNA directed against *Atm*.** Image depicts representative immunoblot for ATM following 4 days of treatment with doxycycline to induce expression of shRNA against *Atm* (shATM) or a non-target control (shNT), using two different shRNA sequences targeting *Atm*. Sequence #1 was highly effective in depleting ATM protein levels compared to non-target controls. Thus, subsequent experiments were performed with shATM #1.

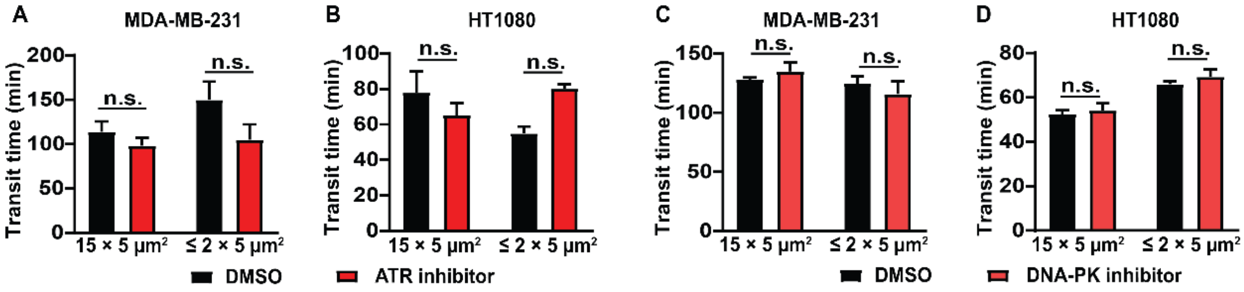


**Supplementary Figure 3: ATR and DNA-PK inhibition do to alter confined migration. (A)** Transit times for MDA-MB-231 cells to cross the ≤ 2 × 5 µm$^2$ constrictions or the 15 × 5 µm$^2$ control channels when treated with ATR inhibitor VE-821 or vehicle control (DMSO). Differences are not statistically significant (n.s.) based on two-way ANOVA with Tukey's multiple comparison test. $N$ = 3 for each group, representing the means of three independent experiments; total number of cells analyzed: 231 cells for DMSO and 184 cells for ATR inhibitor for the small constrictions; 191 cells for DMSO and 129 cells for ATR inhibitor for the larger control channels. **(B)** Transit times for HT1080 fibrosarcoma cells to cross the ≤ 2 × 5 µm$^2$ constrictions or the 15 × 5 µm$^2$ control channels when treated with ATR inhibitor VE-821 or vehicle control (DMSO). Differences are not statistically significant (n.s.) based on two-way ANOVA with Tukey's multiple comparison test. $N$ = 3 for each group, representing the means of three independent experiments; total number of cells analyzed: 388 cells for DMSO and 189 cells for ATR inhibitor for the small constrictions; 388 cells for DMSO and 146 cells for ATR inhibitor for the larger control channels. **(C)** Transit times for MDA-MB-231 cells to cross the ≤ 2 × 5 µm$^2$ constrictions or the 15 × 5 µm$^2$ control channels when treated with DNA-PK inhibitor NU7441 or vehicle control (DMSO). Differences are not statistically significant (n.s.) based on two-way ANOVA with Tukey's multiple comparison test. $N$ = 3 for each group, representing the means of three independent experiments; total number of cells analyzed: 176 cells for DMSO and 255 cells for DNA-PK inhibitor for the small constrictions; 88 cells for DMSO and 115 cells for DNA-PK inhibitor for the larger control channels. **(D)** Transit times for HT1080 fibrosarcoma cells to cross the ≤ 2 × 5 µm$^2$ constrictions or the 15 × 5 µm$^2$ control channels when treated with DNA-PK inhibitor NU7441 or vehicle control (DMSO). Differences are not statistically significant (n.s.), based on Two-way ANOVA with Tukey's multiple comparison test; $N$ = 3 for each group, representing the means of three independent experiments; total number of cells analyzed: 261 cells for DMSO and 326 cells for DNA-PK inhibitor for the small constrictions; 245 cells for DMSO and 259 cells for DNA-PK inhibitor for the larger control channels. Error bars in this figure represent mean ± s.e.m. of the mean values of each experiment.

### Supplementary Videos

**Supplementary Video 1:** Representative example of *Atm* wild-type (WT) and *Atm*-Null nucleus labeled with Hoechst 33342 (blue) undergoing nuclear deformation in micropipette aspiration device with channels of $3 \times 5\ \mu m^2$ in size. Scale bar: 5 µm.

**Supplementary Video 2:** Representative example of an MDA-MB-231 breast cancer cell expressing NLS-GFP (green) and treated with 0.1% DMSO during migration through a $1 \times 5\ \mu m^2$ constriction in the microfluidic device. Scale bar: 5 µm.

**Supplementary Video 3:** Representative example of an MDA-MB-231 breast cancer cell expressing NLS-GFP (green) and treated with 10 µM ATM inhibitor KU-55933 during migration through a $1 \times 5\ \mu m^2$ constriction in the microfluidic device. Scale bar: 5 µm.
